# Differentiation of thermal reaction norms between marginal and core populations of a northward expanding parasitoid

**DOI:** 10.1101/2022.04.26.489532

**Authors:** Emilie Delava, Frederic Fleury, Patricia Gibert

## Abstract

Understanding the speed of and the type of mechanisms that species use to adapt to rapid change is a central question in evolutionary biology. Classically, the two mechanisms denoted in the literature that allow individuals to address these environmental changes are either phenotypic plasticity or rapid evolutionary changes. However, phenotypic plasticity itself can evolve rapidly. In this study, we investigated the genetic differentiation between marginal and core populations of a high-trophic level insect, *Leptopilina boulardi*, a *Drosophila* parasitoid, which has exhibited a very rapid progression northward of its geographical range. Several life history traits have been investigated in different populations according to four fluctuating thermal regimes that mimic the thermal conditions in the field. We found that at low developmental temperature, the two northern marginal populations that have to face a colder winter, survive longer than the two core populations. In addition, the northernmost populations exhibit a higher potential fecundity, a higher starvation resistance and a larger amount of energy at low temperatures. These significant genetic differentiations with genotype-by-environment interactions show that a rapid genetic differentiation of the shape of thermal reaction norms is possible when populations have to cope with new environments.

## Introduction

In recent years, important literature has emerged on the mechanisms of adaptation of species to environmental changes in relation to human activities, be it the impact of climate change or the increase of biological invasions (Chown et al. 2007, 2015; Ghalambor et al. 2007; Hulme 2008; Prentis et al. 2008; Engel et al. 2011; Hoffmann and Sgrò 2011; Bock et al. 2015; Sgrò et al. 2016; Estoup et al. 2016; Gibert et al. 2016). The questions addressed in all these studies are how quickly and by what mechanisms species can adapt to a rapid change of the environment that can generate strong new selective pressures. There are typically three non-mutually exclusive scenarios in the literature by which animals may respond to these changes: i) natural populations can migrate to more favorable geographical areas (e.g. (Parmesan and Yohe 2003; Root et al. 2003; Chen et al. 2011), or remain and adapt to the new conditions ii) through phenotypic plasticity (e.g., Richards et al. 2006; Charmantier et al. 2008; Nicotra et al. 2010; Drown et al. 2011) or iii) by selection for genotypes adapted to the new environmental conditions (evolution) (e.g., (Hoffmann and Sgrò 2011; Urbanski et al. 2012; Hamilton et al. 2015). Plastic responses, i.e. the ability of a genotype to express a different phenotype in various environments, are generally considered to allow rapid responses in a single generation, while evolutionary responses are slower because they require changes in the genetic make-up of populations. However, these two scenarios are not exclusive since plasticity by itself can evolve ((Gotthard et al. 1995; Ghalambor et al. 2007; Scoville and Pfrender 2010) and thus can show greater or lesser variability between populations of the same species (e.g., (Delpuech et al. 1995; Morin et al. 1999; Ayrinhac et al. 2004; Ris et al. 2004; Lind and Johansson 2011; Frei et al. 2014; Pereira et al. 2018). Theoretical studies have suggested that for species continuously distributed in space along an environmental gradient, greater plasticity is expected in marginal populations (Chevin and Lande 2011). How rapidly such an evolution can occur is difficult to assess empirically and requires a situation with a recent and well described range expansion. A comparison of phenotypic plasticity between populations from the core and the marginal recently colonized should allow estimation of this rate of evolution of phenotypic plasticity. In plants that can be easily cloned and that are directly affected by environmental variations because of their inability to move, this type of study comparing geographic and altitudinal ecotypes exists (e.g. (Nicotra et al. 2010). In insects, except for some iconic (e.g. butterflies) or harmful (e.g. mosquitoes) species, long-term monitoring is rare. Moreover, studying phenotypic plasticity requires the possibility of conducting laboratory experiments that control the environmental variation, a situation that is not possible in all organisms.

*Leptopilina boulardi* (Hymenoptera: Eucoilidae) is a larval endoparasitoid that uses *Drosophila* as a host, mainly *D. melanogaster* and *D. simulans*, to achieve its larval development until it is a free-living adult. The geographical distribution of *L. boulardi* ranges from tropical to subtropical areas in Africa and America, including areas with a Mediterranean climate (Hertlein 1986; Allemand et al. 2002). In Europe, *L. boulardi* has been reported all around the Mediterranean basin in Spain, southern France, North Africa and the Middle East ((Barbotin et al. 1979; Nordlander 1980; Allemand et al. 2002; Fleury et al. 2009). (Delava et al. 2014) showed that in the southeast of France, *L. boulardi* is moving very rapidly northwards, with an average rate of range expansion of 170 km in 19 years (1993-2011), exceeding previously observed rates for small insects. This range expansion is not limited by host availability and could be related to climate warming since this area was characterized by an annual warming of temperature with a northward displacement of isoclimatic lines by approximately 100-130 km over 33 years (Lelièvre et al. 2011). However, during the same period, winters have become colder, with an average temperature decrease of -1.58°C that may imply the adaptation of the species to new local conditions.

In this paper we study how and how quickly latitudinal temperature variations can change the shape of reaction norms. We thus compared thermal reaction norms of various life history traits between four populations, two populations 150 km apart located in the central area of *L. boulardi* and two populations about 20 km apart located in the recently colonized northern area of *L. boulardi* (marginal populations) range. We expect a genetic differentiation of fitness-related traits between populations from marginal and core areas in response to these different thermal selective pressures, with possible genotype-by-environment interactions.

## Materials and methods

### BIOLOGICAL MODEL, COLLECTION SITES AND LABORATORY REARING

*Leptopilina boulardi* is a solitary (only one offspring can survive per host whatever the number of eggs deposited) endoparasitoids that attacks first and second stages of Drosophila larvae (Carton et al. 1986). In many cases, the issue of parasitism is the emergence of a parasitoid but there can also be a precocious parasite death induced by the immune response of the host (called encapsulation) leading to the emergence of a Drosophila. In some cases, also, a physiological inadequacy can lead to the death of both partners (Fleury et al. 2009). In *L. boulardi*, females can be infected by a maternally transmitted DNA virus known as LbFV (Leptopilina boulardi filamentous virus), frequently leading to a situation of superparasitism (deposition of supernumerary eggs into previously parasitized larvae) (Varaldi et al. 2003). Previous work showed Lbfv has no detectable effect on female survival, but has a slightly positive effect on egg load and a weakly negative effect on tibia length and developmental rate (Varaldi et al. 2005).

Parasitoid populations were collected from orchards using several banana bait traps (at least 7 traps per population) placed in four sampling localities in southeastern France (Supplementary S1). Two sampling sites, Eyguières (S1, Latitude 43°41’N) and Gotheron (S2, Latitude 44°58’N), were located in the south of the Rhône-Saône Valley, where a Mediterranean climate prevails and where *L. boulardi* has been well established for many years (core area). The two other localities, Sainte-Foy-lès-Lyon (N1, Latitude 45°44’N) and Saint-Maurice-de-Beynost (N2, Latitude 45°49’N), were located in area where *L. boulardi* was absent in the 1990s, but that was progressively colonized by this species during the last twenty years (marginal area) (Delava et al. 2014). The most southern central population and the most northern marginal population are separated by approximately 240 km.

Parasitoids sampled in the 4 localities were used to established 4 mass populations of *L. boulardi* that were reared on a standard laboratory strain (more than 250 generations in the lab) of *Drosophila melanogaster* in a controlled environment at 25 ± 0.3°C, 70% RH (Sanyo MLR-351H climatic chamber), with a photoperiod of 14L:10D. The experiments were conducted after approximately 10 generations in the laboratory.

### TEMPERATURE TREATMENTS DURING DEVELOPMENT

*L. boulardi* shows a thermal specialization with an optimum around 25 °C, and a fall of its performance when the temperature deviates by more than 2 °C from this optimum (Fleury et al. 2009; Moiroux et al. 2013). It is an over-wintering species with a facultative larval diapause, almost 100% of the larvae entered into diapause at 17.5 °C (Claret and Carton 1980). Because of this diapause, developmental plasticity has never been investigated in this species, but it has been recently shown that the use of a fluctuating thermal regime allows the larval diapause to be overcome (Delava et al. 2016). In this paper, we have thus simulated the daily temperature fluctuation as closely as possible to the natural thermal regime. Towards that aim, temperature was recorded every hour (EasyLog USB data logger) in seven localities of the Rhône-Saône valley from latitude 44°58’N to latitude 46°24’N over 62 days (from July 1 to August 31, 2010), when the abundance of *L. boulardi* is high. For each hour of the day, the median of the 62 days was calculated and used to create a fluctuating thermal regime that showed an average temperature of 22.7°C. This thermal regime was then modified by increasing the temperatures by + 1.5°C, - 1.5°C and - 3°C to obtain 4 fluctuating thermal regimes with the same amplitudes but with different average temperature: 19.7°C, 21.2°C, 22.7°C and 24.2°C (Supplementary S2). In the following text, fluctuating thermal regimes will be named using these average values.

The experiment was conducted simultaneously in 4 identical climatic chambers (Sanyo MLR-351H) in which the temperature was controlled according to the tested thermal regime with a precision of 0.1°C. In the 4 chambers, the temperature varied every two hours, while the photoperiod and humidity were the same (12L: 12D, 70% RH), and the temperature was recorded with a datalogger (EasyLog USB).

### EXPERIMENTAL DESIGN

One hundred twenty-five *D. melanogaster* eggs were deposited into rearing vials (15 replicates per thermal regime and per population, 240 vials in total) (see Fig. 1). Vials were then randomly assigned to the 4 incubators (60 vials per incubator in total). After 24 h (to allow eggs to hatch), a single female parasitoid was introduced into each vial and removed 24 h later.

**Figure 1.**
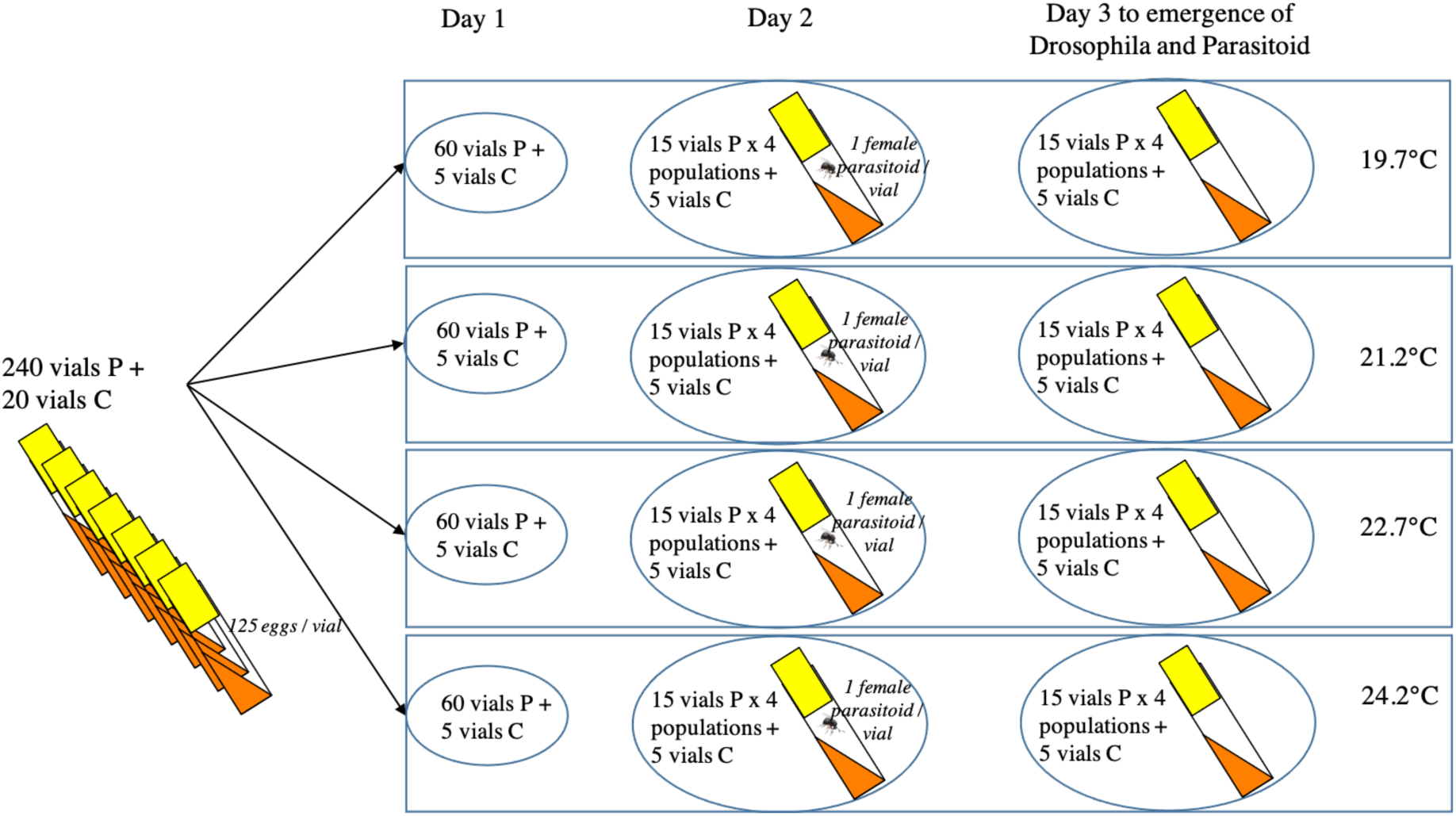
Schematic representation of the experimental protocol used: 125 D. melanogaster are deposited in each tube in which a female *L. boulardi* is placed for 24H. 15 replicates are made for each population and each thermal regime. For each condition, 5 control tubes without parasitoids are used to evaluate the quality of Drosophila development.

All females were kept to check the presence of LbFV viral particles by PCR using the protocol described by (Patot et al. 2009). We confirmed previous results found in (Patot et al. 2010), i.e. no parasitoid from N1 and N2 (populations from marginal region) were infected by the virus LbFV, while most parasitoids from S2 and S1 (populations from core region) were infected by LbFV (78% at S2 and 94% at S1 were infected).

For each thermal regime, five control vials (20 in total) with only the 125 *D. melanogaster* eggs but without *L. boulardi* were used to assess the quality of the development of *Drosophila* larvae in the absence of parasitoids.

All vials were maintained under the 4 fluctuating thermal regimes until the insect’s complete development. Adult females used to measure adult life history traits, such as lifespan and fecundity, were kept under the same thermal conditions as larval development.

### TRAITS RELATED TO THE DEVELOPMENT OF THE PARASITOID

The insects’ emergence was checked daily, and all adults (*Drosophila* and then parasitoids) were counted in each vial. To estimate the host immune response, adult *Drosophila* were dissected under a microscope by crushing the entire individual between two glass slides, and the number of flies containing encapsulated parasitoids egg (the immune response of *Drosophila* host) was recorded. Since the number of capsules was negligible (less than 1%) and was not significantly different by the thermal regimes (GLM, χ^2^_(3,221)_ = 335, *P* = 0.47) and populations (χ^2^_(3,218)_ = 324, *P* = 0.15), we did not include this parameter in further estimates of parasitoid development.

Using the number of emerging Drosophila and parasitoids counted daily in each vial, two parameters related to the host-parasitoid interaction were calculated.

#### Host infestation rate (IR)

This parameter measures the number of *Drosophila* parasitized by wasps no matter the issue of parasitoids development. IR was calculated as IR_i_ = (nc – n_di_) / nc, where nc is the average number of *Drosophila* emerging from control vials and n_di_ is the number of *Drosophila* that emerged from each test vial.

#### Success of parasitism (SP)

This trait estimates the percentage of parasitized hosts that give rise to adult parasitoids (larval survivorship of the wasps). SP was calculated for each female’s offspring as follows: SP_i_ = n_pi_ / (125 – (n_di_ / s)), where n_pi_ is the total number of parasitoids produced in each vial, 125 is the total number of *Drosophila* hosts eggs in each vial, n_di_ is the total number of adult flies emerging from each vial (and thus that escaped from parasitism) and s is the survival rate of flies emerging from control vials (s = nc / 125).

We also measured the ***Parasitoid development time***, i.e., the development duration of the parasitoid from the day of deposit of *Drosophila* eggs in vials to the emergence of an adult parasitoid.

### ADULT PARASITOID PHENOTYPIC TRAITS

#### Potential fecundity

Egg-load was measured on 5-day-old honey-fed females (20 per population and per developmental thermal regime, 320 in total). Parasitoid females were individually dissected in a drop of Ringer’s solution under a binocular microscope (BBT Krauss). One of the two ovaries was placed under a microscope (Axio Imager, AxioCam, Software Axiovision LE; Zeiss, Thornwood, NY, USA) and was photographed. The eggs were counted on each photograph using ImageJ. The total fecundity of the individual was estimated as being twice the number of eggs contained in one ovary.

#### Size

The size of females tested for fecundity was determined by measuring the length of their right tibia using the same microscope. Tibia length was shown to be a good proxy of individual size in parasitoids (Cronin and Strong 1996; Nicol et al. 1999), and individual size is positively correlated with most life-history traits (Godfray 1994).

#### Starvation resistance

After emergence, some parasitoid females were placed in vials without a host and food but with sterile moistened cotton to measured their lifespan (10 females per vial and 4 vials per population and per thermal regime, 640 females in total). Cotton was moistened daily to provide humidity. Dead parasitoids were counted twice a day, every morning and evening (at 9:00 a.m. and 6:00 p.m.).

### ADULT PARASITOID ENERGETIC RESOURCES

A part of some newly emerged females was stored at -20°C to measure their energetic reserves. Proteins, lipids, sugars and glycogen content were measured according to the protocol of (Foray et al. 2012). Since *L. boulardi* is a small insect (approximately 2 mm), preliminary analyses were conducted to test the sensitivity of the method for all metabolic compartments. On the basis of this preliminary test, we decided to pool 8 individuals to perform extraction on 10 repetitions per population and per thermal regime (160 samples in total). The value obtained in each energy compartment for a pool of 8 individuals was divided by 8 to obtain an average amount per individual. Because the quantities of energy reserves are correlated with the individual’s size, we divided all quantities (for each compartment) by the average size measured from 20 females per population and thermal regime to take into account this size effect. Then, the total energy amount in J/mg available at emergence was assessed using the following conversion factors: 16.74 J/mg carbohydrates and proteins and 37.65 J/mg lipids (Rivero and Casas 1999). Finally, the amounts of proteins, lipids, sugars and glycogen (in joules) were divided by the total amount of energy in order to obtain the percentage of each compartment. Because the amount of free sugars and glycogen was very low (2.5% and 8.0%, respectively), they are not presented in this study.

### STATISTICAL ANALYSES

All analyses were performed with the R software (version 2.14.1) (R Development Core Team, (2011). Vials with no parasitoid emergence were excluded from the analysis. Except for the number of capsules and adult survival, all life history traits were analyzed using linear models after adequate transformation to comply with the assumptions of normality and homoscedasticity when necessary (log transformation for 1-IR and arcsine square-root transformation for SP). A generalized linear model was used to analyze the number of capsules with a quasi-Poisson error (log link function) to take into account the over-dispersion of the trait. Survival was analyzed by means of a Weibull distribution using the function “survreg” from the R package “survival”. We included vials as a random effect using the “frailty” function, but since it did not improve the model, this effect was removed. In all models, the population (qualitative variables) and the developmental temperature regime (quantitative variable) were tested as fixed effects. We have also included a quadratic or cubic effect of developmental temperature to better fit the shape of the performance curve when appropriate. Initial models were simplified by stepwise regression while minimizing Akaike’s information criterion (AIC). The results include only the best models selected by the lowest AIC criterion (Johnson and Omland 2004).

A linear model was also performed to check whether the potential fecundity was correlated with body size (estimated by the length of the tibia). Since correlations between these two variables were found non-significant, the size was not included in the model explaining fecundity.

For all models, comparisons between populations and thermal regimes were then possible by changing the reference population and thermal regime in the contrast matrix.

## Results

### TRAITS RELATED TO THE DEVELOPMENT OF THE PARASITOID

#### Host infestation Rate (IR) and Success of Parasitism (SP)

No significant differences were observed for IR, no matter the developmental thermal regime or the population. SP exhibited a significant interaction between the population effect and the cubic effect of developmental thermal regime (*F* = 5.23, *P* = 0.0018) (Fig. 2A), with a lower SP value for the population of S2 at 24.2°C.

**Figure 2.**
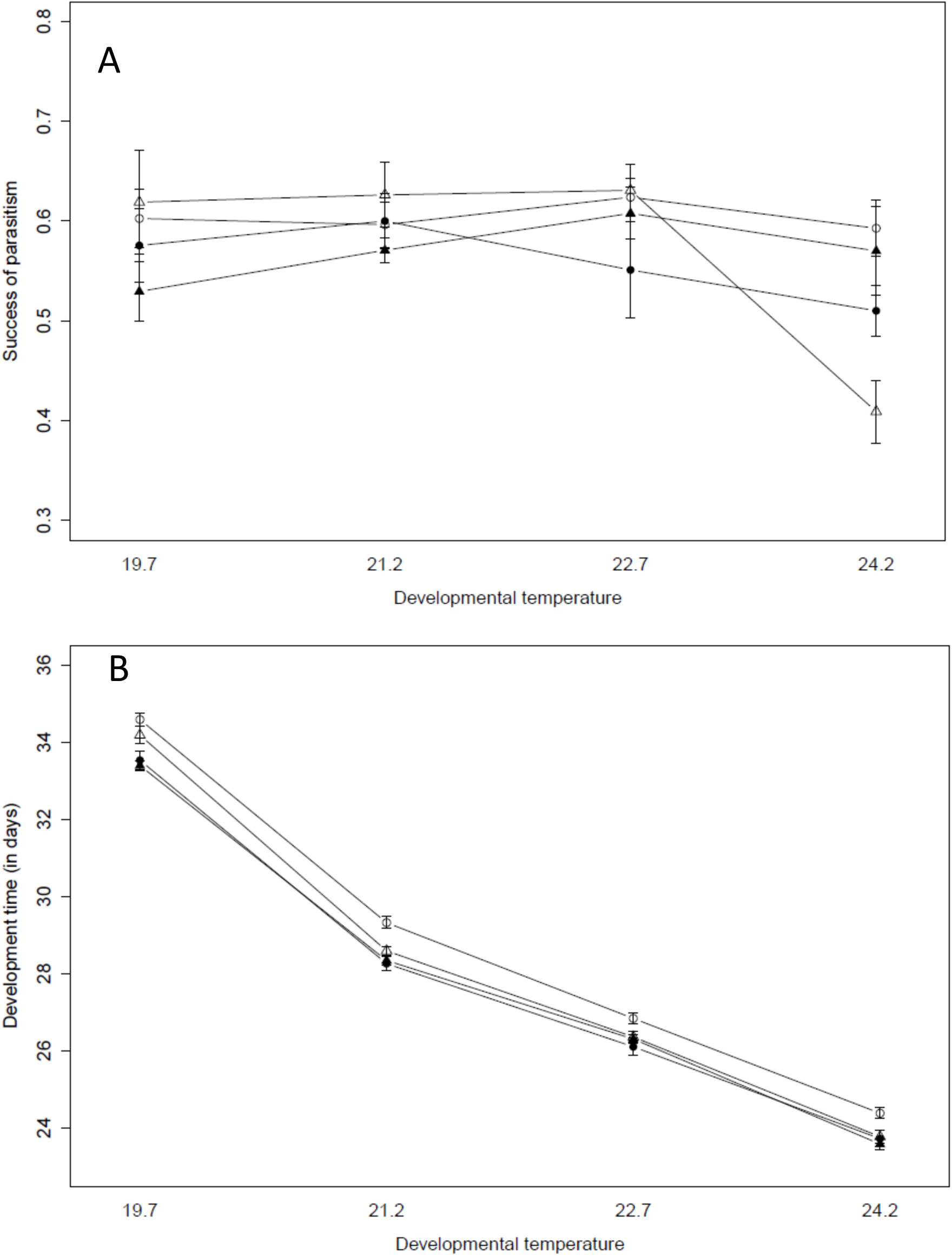
Reaction norms of A) success of parasitism and B) development time across the four fluctuating developmental thermal regime. Each point represents the mean value per population (± SE). Open symbols represent the two core populations (triangle: S2 and circle: S1) and symbols in black represent the two marginal populations (triangle: N2 and circle: N1).

#### Development time

Development time significantly varied with both developmental thermal regime and populations, but no significant interaction was observed. Development time decreased when the average temperature of the developmental thermal regime increased (*F* = 7867.28, *P* < 0.001) (Fig. 2B) as classically observed for insect species. The development duration for the complete eggs to adult development was approximately 24 days at 24.2°C and increased until 34 days at 19.7°C. Development time also differed significantly between populations (*F* = 27.10, *P* < 0.001). The southern population, S1, developed significantly slower than the three other populations (28.87 days±3.84 for S1 versus 27.91 days on average for the three other populations).

### ADULT PARASITOID LIFE-HISTORY TRAITS

#### Potential fecundity

For the four studied populations, female egg-load showed no significant differences between the developmental thermal regimes (*F* = 0.0034, *P* = 0.95). We thus pooled the data of these thermal regimes and compared them between populations. A strong population effect was observed (*F* = 21.62, *P* < 0.001), with females from the northern population, N2, showing a significantly higher egg-load (158 ±21.9) than the three other populations (N1, S2 and S1) (133 ± 30.7 on average) (Supplementary S3).

#### Size

A significant effect of both population and developmental thermal regime was found on size, with no significant interaction. Surprisingly, and contrary to what is generally found in insects, size estimated by the tibia length significantly increased with developmental temperature (*F* = 7.41, *P* = 0.0017) (Fig. 3A). We also found that females from the southern population, S1, were significantly bigger (301.61 ±9.94 μm) than those from N2 and S2 (297.62 ±11.47 μm and 294.77 ±11.29 μm respectively).

**Figure 3.**
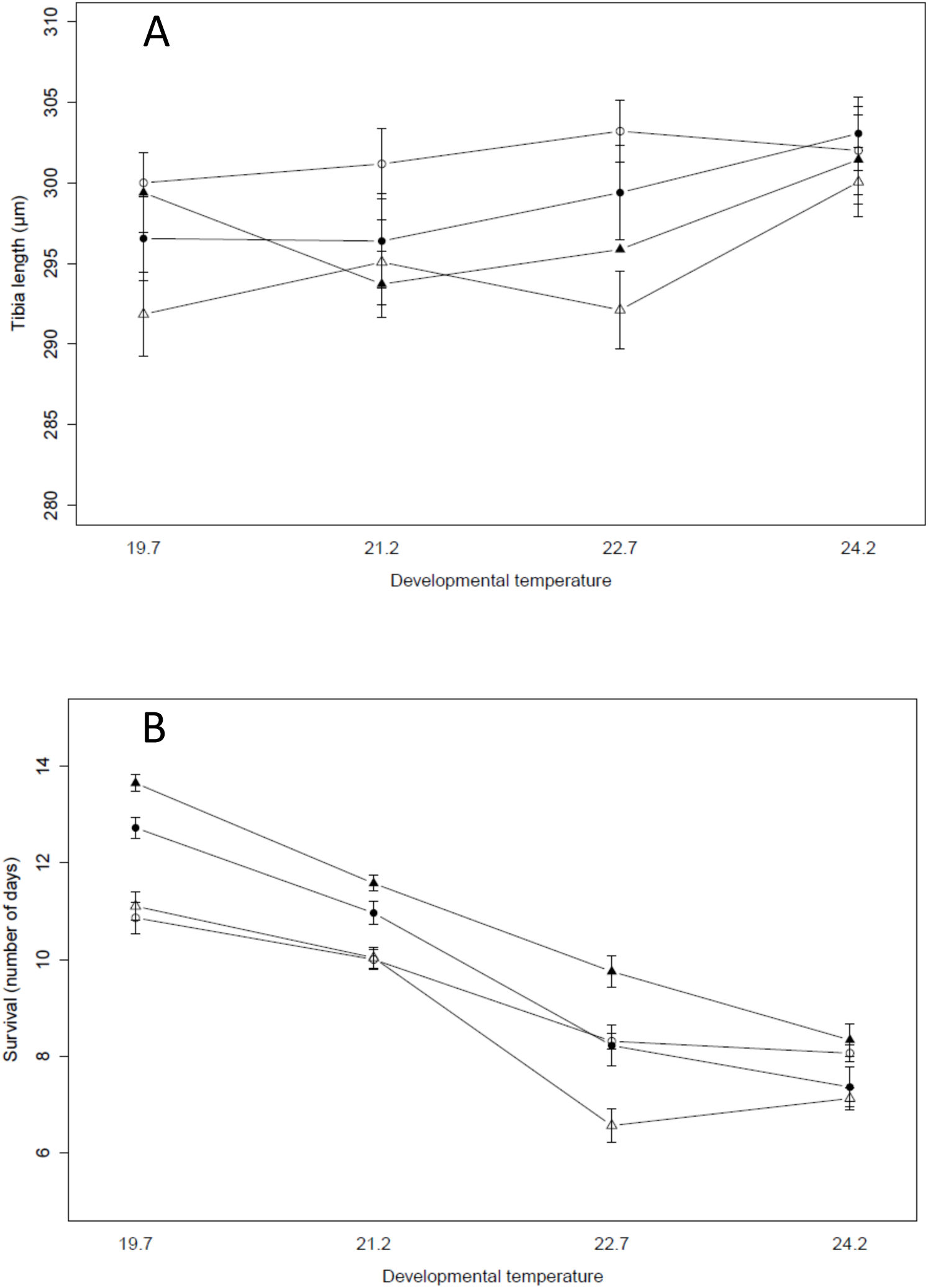
Tibia length A) and starvation resistance B) (mean±SE) reaction norms across four developmental thermal regimes. Open symbols represent the two core populations (triangle: S2 and circle: S1) and symbols in black represent the two marginal populations (triangle: N2 and circle: N1).

#### Starvation resistance

A significant interaction between developmental thermal regime and population (Dev = 9.44, *P* = 0.024) was observed for adult survival. As it is observed for many insects, we observed a decrease of survival when temperature increases for all populations (Dev = 414.13, *P* < 0.001). However, we found that at the lower thermal regime (19.7°C and 21.2°C), parasitoids from the northern marginal region (N2 and N1) lived significantly longer than parasitoids from southern populations (S2 and S1), whereas at 24.2°C no significant differences were observed between the four populations (Fig. 3B).

### ADULT PARASITOID ENERGETIC RESOURCES

The developmental thermal regime significantly influenced the total amount of energy, which decreased when temperatures become higher (*F* = 75.99, *P* < 0.001), with a significant quadratic and cubic effect (*F* = 12.44, *P* < 0.001 and *F* = 15.53, *P* < 0.001 respectively).

Females developed at the two lower temperatures of 19.7°C and 21.2°C had significantly more energy than females developed at 22.7°C and 24.2°C (Fig. 4). In addition, there was a significant difference in the total amount of energy between populations (*F* = 5.57, *P* = 0.0012). Females from N2 had more energy than females from the three other locations, especially at 19.7°C and 21.2°C, even if the interaction between population and developmental thermal regime was not significant (interaction with linear effect, *F* = 1.83, *P* = 0.14; interaction with quadratic effect, *F* = 2.00, *P* = 0.12; interaction with cubic effect, *F* = 2.63, *P* = 0.053).

**Figure 4.**
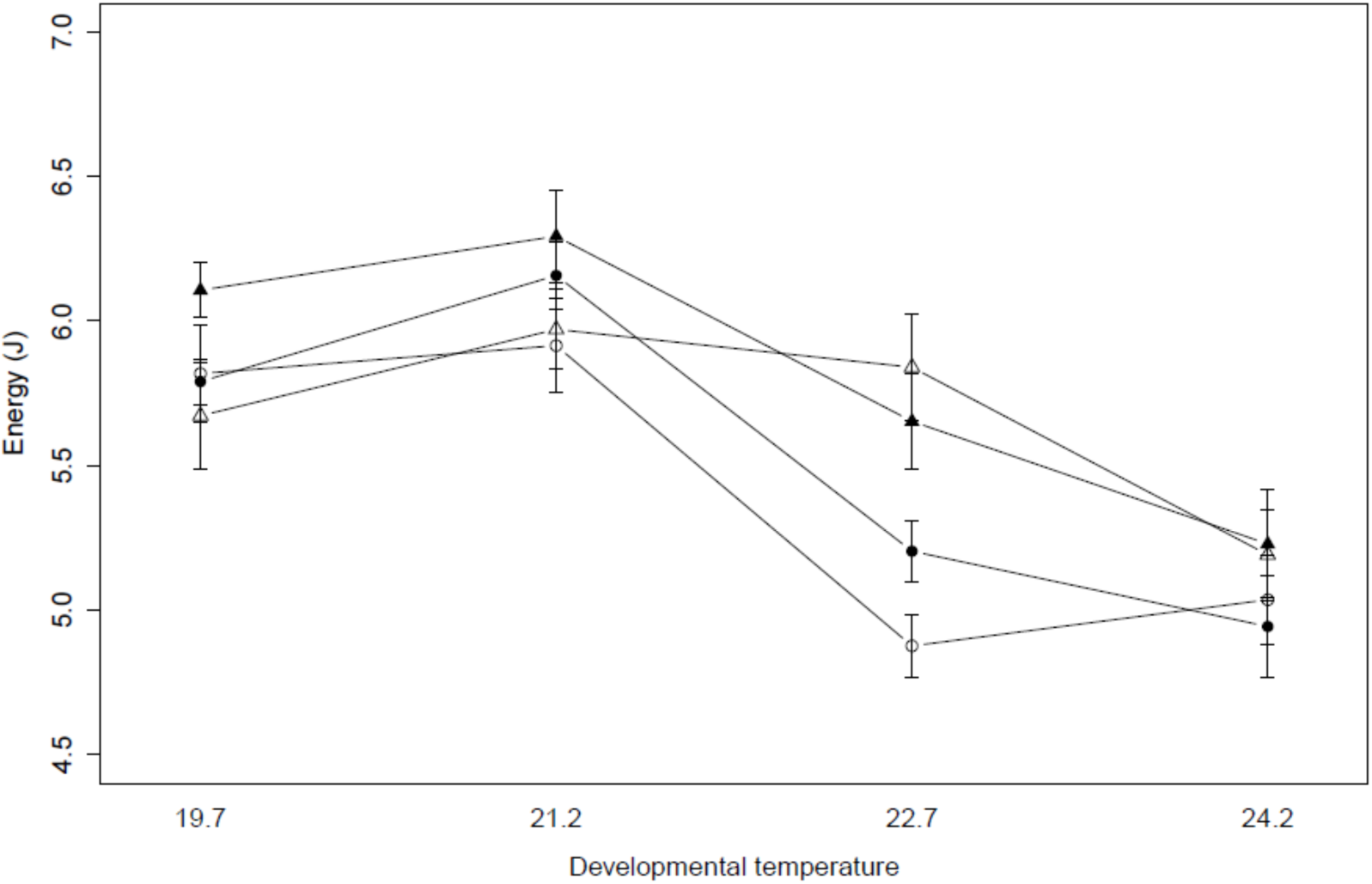
Energy content (mean±SE) reaction norms across four developmental thermal regimes. Open symbols represent the two core populations (triangle: S2 and circle: S1) and symbols in black represent the two marginal populations (triangle: N2 and circle: N1).

Whereas no developmental thermal regime by population interaction was detected for the total amount of energy, a genotype-by-environment interaction was significant for both protein and lipid rates, but with an opposite trend. For the protein rate, a highly significant developmental thermal regime by population interaction (*F* = 22.38, *P* < 0.001) resulted in a lower protein content in northern marginal populations (N2 and N1) at higher temperatures, whereas all populations were similar at low temperatures (Fig. 5A). In contrast, for the lipid rate, the significant interaction between the variables population and developmental thermal regime (*F* = 13.48, *P* < 0.001) was due to a higher rate for northern marginal populations at the highest temperature of 24.2°C, whereas small differences among populations were observed at low temperatures (Fig. 5B).

**Figure 5.**
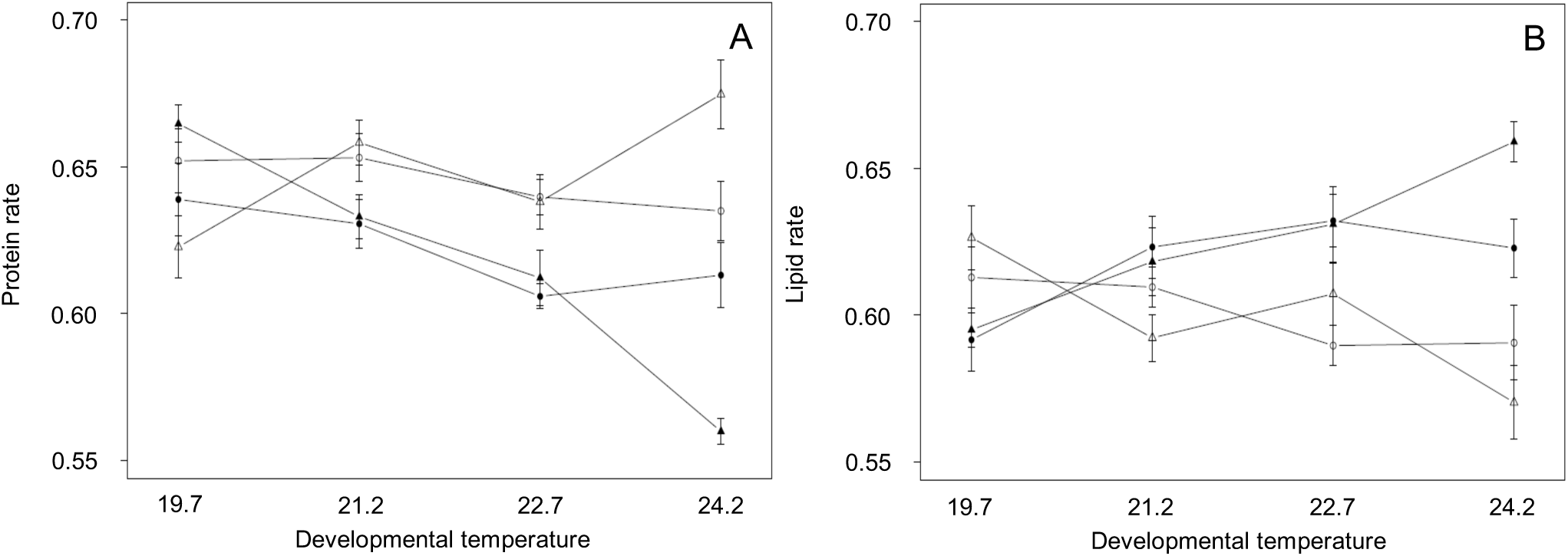
A) protein rate and B) lipid rate (mean±SE) reaction norms across four developmental temperatures. Open symbols represent the two core populations (triangle: S2 and circle: S1) and symbols in black represent the two marginal populations (triangle: N2 and circle: N1).

## Discussion

The objective of this paper was to describe the reaction norms of an insect whose range has recently been extended in relation to global warming. Several traits were thus studied under four different thermal regimes and for four of the central and marginal range populations. The use of fluctuating temperatures allowed us to circumvent the parasitoid diapause (Delava et al. 2016), which is induced by relatively high temperatures, since 50% of parasitoids enter into diapause under a constant 20°C (Claret and Carton 1980). Our study thus constitutes the first investigation of the phenotypic plasticity of *L. boulardi* under this range of temperatures.

### DIFFERENTIATION OF THERMAL REACTION NORMS OF LIFE HISTORY TRAITS

We found a significant differentiation between populations for various traits. The southern population, S1, develops faster and is bigger than the other populations, while the northern population, N2, has the highest fecundity.

According to the temperature-size rule, the typical pattern in ectotherms is that body size is negatively correlated with developmental temperature (Atkinson 1994; Angilletta and Dunham 2003). Our results depart from this rule since we found that the tibia length (the proxy of body size) regularly increases when the fluctuating developmental thermal regime increases. To our knowledge, this is the first time that such a result is highlighted in ectotherms. Another intriguing result is observed for potential fecundity, which is not affected by fluctuating developmental thermal regime, in contrast to the concave reaction norms that are generally observed using constant temperatures in insects (e.g., (Delpuech et al. 1995) on *Drosophila melanogaster* or (Ris et al. 2004) on *Leptopilina heterotoma*). Our results suggest therefore that using fluctuating temperatures can buffer the impact of developmental temperature at least for these traits and questions the generalization of the results using only constant temperatures.

A significant population-by-temperature interaction reflecting a modification of the shape of the thermal reaction norms between populations is found for two traits only: the success of parasitism and starvation resistance. Clearly, starvation resistance is the trait for which this effect is the most interesting. Indeed, for the two coldest thermal regimes (fluctuating developmental thermal regime of 19.7°C and 21.2°C), survival is significantly higher for the two marginal northern populations (N2 and N1) than for the core populations (S2 and S1), while in the warmer regime, 24.2°C, there is no difference between populations. Since in the field, the maximal and minimal temperatures are lower in the marginal than in the core region (Delava et al. 2014), our results suggest a local adaptation of marginal populations that survive better at low temperatures than at higher temperatures compared to the core populations.

If the thermal gradient of the Rhone valley alone were responsible for the differences in reaction norms between populations, one would expect these differences to be correlated with the geographical distance between the populations. However, we observe that for the two populations in the core area, which are very far apart, the shape of the reaction norms is similar and very different from the shape of the reaction norms of the two populations in the marginal area. Instead, our results suggest an adaptation of the populations to particular environmental conditions in the marginal zone. In this northern zone, winter remains very cold and an average temperature increase was only observed in spring and fall (Delava et al. 2014). The observed change in the shape of the response norms for starvation could therefore be related to the ability of populations to persist in the marginal zone during the cold season.

### ENERGETIC CONTENT ANALYSIS

The overall energy content decreased with the fluctuating developmental thermal regime and varied among populations. In fact, the northern population (N2) had significantly more energy than the three other populations especially at the lowest developmental temperature. In parasitoids, the nutritional resources are provided by the host and are stored by the parasitoid during ontogeny to constitute the only energy reserves available at emergence (Rivero and Casas 1999; Pelosse et al. 2007). *L. boulardi* is a pro-ovigenic parasitoid (its egg stock is complete and mature at emergence), which means that the total energy at emergence will be exclusively allocated to maintenance and locomotion. Thus, the greater amount of energy of N2 at 19.7°C can probably explain its better starvation resistance in a cold environment.

Because free sugars and glycogen represented negligible amounts of energy compared to proteins and lipids, we have focused on these two latter energy compartments. A significant population-by-temperature interaction was found for these two traits, with similar values between populations in the low fluctuating developmental thermal regime but a significant differentiation between populations in the high fluctuating developmental thermal regimes. However, opposite tendencies were observed since the marginal populations (N2 and N1) have a lower protein content but a higher lipid content than the core populations (S2 and S1). These high fluctuating developmental thermal regimes were characterized by temperatures above 30°C, which should constitute a particularly stressful environment, especially for the marginal northern populations. One of the specific traits of the parasitoids is that they are unable to synthesize lipids during adult life (Visser and Ellers 2008); as a consequence, the fat reserves accumulated during development should presumably provide a significant advantage for resisting extreme environments.

### Possible confounding effect with LbFV

In the present study, we also confirm the initial observations of (Patot et al. 2010) that populations in the central region (S2 and S1) are infected with LbFV, while those newly established in the northern areas (N2 and N1) are free of infection. LbFV is known to influence the life history traits of *L. boulardi* (Varaldi et al. 2005), and this effect is potentially confounded by geographic variation in populations. However, LbFV infection is known to cause a slowing of host development (Varaldi et al. 2005), yet we observe significantly shorter development for the two populations in the marginal region (N2 and N1 uninfected) compared to that of S1 (the southernmost population in cline, almost entirely infected with LbFV). Similarly, for fecundity, infected individuals are supposed to have a higher egg number than uninfected individuals (Varaldi et al. 2005) whereas it is the N2 population (the most northern population of cline, uninfected by LbFV) that exhibited the highest potential fecundity, reinforcing the hypothesis of a local adaptation of this population. These results suggest that even if we cannot totally exclude an effect of the presence of LbFV on the life history traits of the studied populations, this effect is not sufficient to hide the genetic differentiation between populations.

## Conclusion

Newly established marginal populations of *L. boulardi* significantly differed from core populations for several life history traits. The northern population (N2) is particularly well differentiated, with a higher starvation resistance, a higher potential fecundity and overall a higher energy content. In this population, *L. boulardi* was observed for the first time five years before we sampled the individuals for this study, showing that a rapid differentiation of thermal reaction norms is possible and thus that the evolution of phenotypic plasticity can be fast. This differentiation could be the result of natural selection but also due to random genetic drift that can be frequent in marginal populations often characterized by small size. In this area, it has been shown that an increase of temperature in spring and autumn probably allowed the northward displacement of *L. boulardi* but also that the coldest temperature during winter could constitute an important selective pressure that could explain the local adaptation of populations. To study the genetic structure of populations, the intensity of gene flows between populations and the dispersive capacity of *L. boulardi* more finely, investigations of population genetics using neutral molecular markers (e.g., RadSeq) are required.

## Acknowledgement

This work is part of the ANR CLIMEVOL project funded by the Agence Nationale de la Recherche. We thank the INRA center of Gotheron, and the landowners who allowed us to collect insects in their orchards. We are also grateful to Lucas Leger and Isabelle Amat who advised us on statistical analyzes. Finally, we would like to pay homage to our dear colleague Roland Allemand, who greatly contributed to the parasitoid sampling, the development of the protocol and provided valuable advices during all this study, but sadly passed away in early 2013.

