## Supplementary S1 for "Differentiation of thermal reaction norms between marginal and core populations of a northward expanding parasitoid"

Supplementary S1 - Location of the 4 localities where parasitoids were sampled: Eyguières (S1, Latitude 43°41’N), Gotheron (S2, Latitude 44°58’N), Sainte-Foy-lès-Lyon (N1, Latitude 45°44’N), Saint-Maurice-de-Beynost (N2, Latitude 45°49’N).

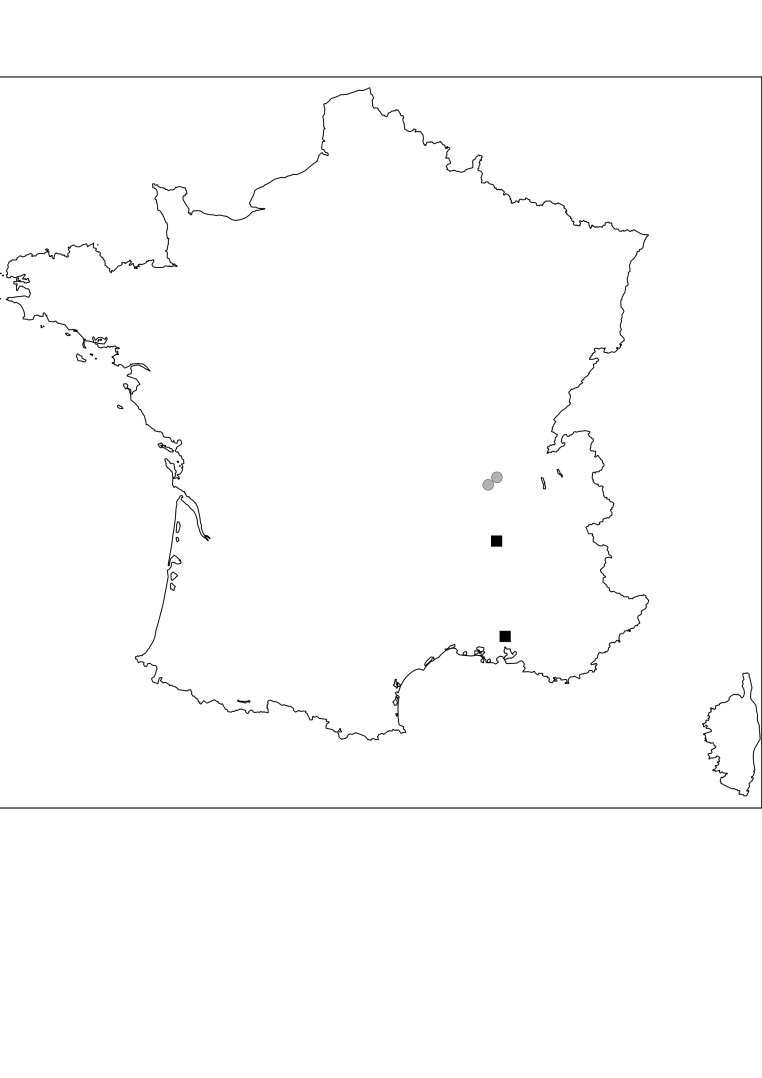

**SF**

**SM**

**GO**

**EY**
