## Supplementary figures and images for "Differentiation of thermal reaction norms between marginal and core populations of a northward expanding parasitoid"

### Supplementary S2

Supplementary S2 - Fluctuating thermal regimes used as environmental conditions.

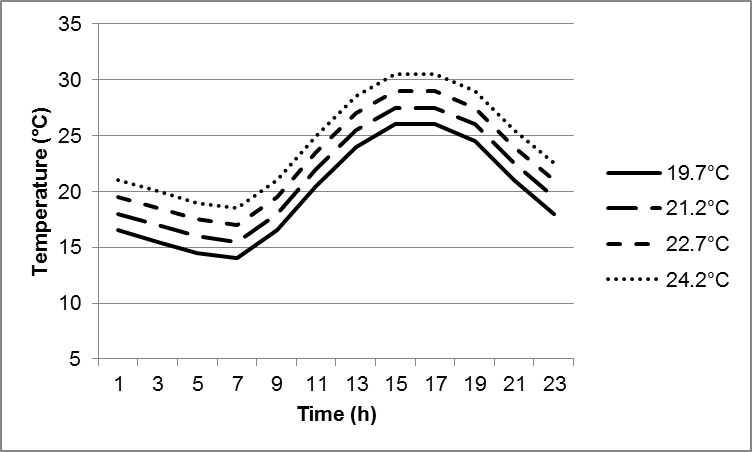
