## Supplementary S3 for "Differentiation of thermal reaction norms between marginal and core populations of a northward expanding parasitoid"

Supplementary S3 Potential fecundity observed for the 4 populations (data not being significantly different between thermal regimes were pooled).

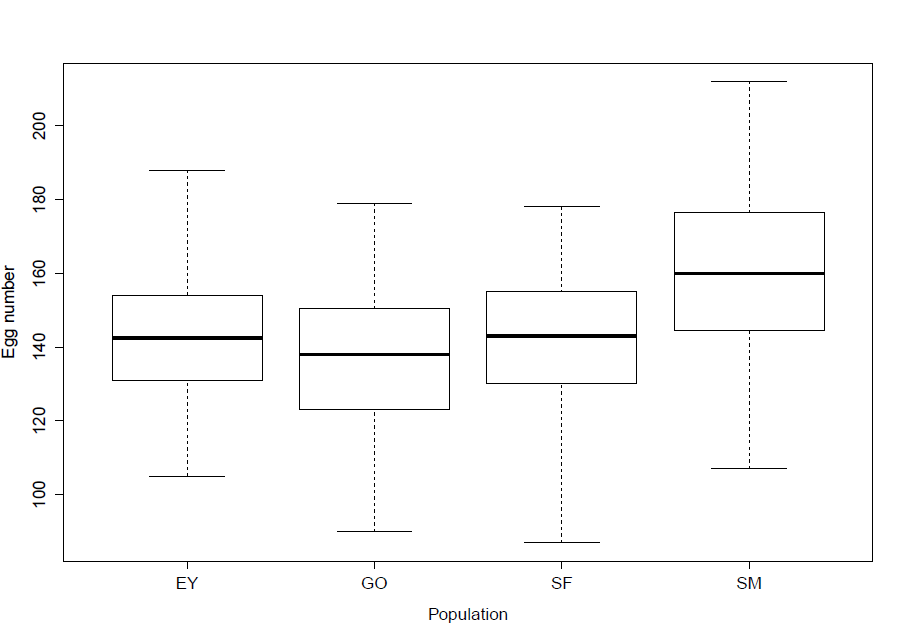
